## Supplementary Methods and Figures for "SurVIndel2: improving CNVs calling from next-generation sequencing using novel hidden information"

#### Determining number of repeat units deleted

The Human Genome SV Consortium (HGSVC) has created a comprehensive catalogue [7] (named HGSVC2) of the SVs in 35 samples: 34 samples from the 1000g project (including HG00512), plus HG002 (alternatively called NA24385). Using these two catalogues, we investigate whether most CNVs in the human genome delete or duplicate whole repeat units. We determined the number of tandem repeat units deleted by the examined deletions as follows. We downloaded TRF [2] annotations for hg38 from the UCSC Genome Browser. For each deletion, we found the tandem repeat that contains the deletion; when we found multiple such repeats, we selected the one with the shortest repeat unit. The number of deleted copies was simply obtained by dividing the length of the deletion by the length of the repeat unit.

#### Building accurate alternative alleles

By building an accurate alternative allele for each CNV and aligning the reads to it, we can precisely identify reads that are supposed to support it. We downloaded PacBio HiFi reads for both HG00512 (SRR13606079 to SRR13606084) and HG002 (SRR10382244 to SRR10382249), and aligned the reads to hg38 using minimap2 [14].

First, we selected a subset of CNVs so that no two CNVs were within 4000 bp of each other. Then, for a given CNV $C$, let $s$ be the start of $C$ minus 2000 bp, and $e$ be the end of $C$ plus 2000 bp. For each HiFi read overlapping $C$, we extract the portion overlapping the region from $s$ to $e$. In case of heterozygosity, we want to identify the subset of reads supporting $C$. In order to accomplish this, for each subread, we compute two numbers $d$ and $i$: the sum of the lengths of all deletions and insertions, respectively ($\geq$10 bp in order to exclude most sequencing errors). Then, we cluster the reads according to $d$ and $i$ using DBSCAN [9]. Each cluster represents an allele. In order to find the cluster that represents the allele with $C$, we find the cluster that minimises the following formula

$$\sum_{r \in S} ( \min_{x \in indels(r)} (size(x) - size(C))^2$$

where $S$ is a cluster of reads and $indels(r)$ is the set of indels in a read $r$. Intuitively, we want to find the cluster with the minimum sum of squared errors, where the error in a read is the difference between the length of the expected CNV and the observed one. Finally, we use the selected reads to assemble the alternative allele using SPOA [28].

Next, we aim at selecting a subset of alleles that are extremely likely to be correct. We apply a series of filters to our set of assembled alleles:

- **No reads**: no reads span the whole region from $s$ to $e$, therefore no alternative allele could be built;
- **No SV**: when aligning the built alternative allele to the reference, no SV $\geq$50 bp could be identified;
- **Multiple SVs**: when aligning the built alternative allele to the reference, multiple SVs $\geq$50 bp could be identified;
- **Incompatible SV**: when aligning the built alternative allele to the reference, an SV could be identified, but the length was very different from the expected CNV (size difference >100 bp);

| | **HG002 DEL** | **HG002 DUP** | **HG00512 DEL** | **HG00512 DUP** |
|---|---|---|---|---|
| No reads | 126 | 106 | 224 | 179 |
| No SV | 467 | 232 | 417 | 214 |
| Multiple SVs | 111 | 217 | 119 | 202 |
| Incompatible SV | 288 | 576 | 262 | 551 |
| No good LR support | 845 | 734 | 729 | 833 |
| Other | 45 | 58 | 41 | 44 |
| High quality | 6150 | 4796 | 6148 | 4585 |

Table 1: Each putative alternative allele undergoes a strict filtering process, in order to remove potentially incorrect sequences.

- **No long-read support**: when realigning the HiFi reads to the alternative allele using minimap2, we expect to find at least 3 reads that (i) cover the whole alternative allele and (ii) they have no large indels (>10bp) and a low error rate (<0.1%). Furthermore, we use these reads to call phased variants using PEPPER-Margin-DeepVariant [23]. If the allele is correctly built, we expect one of the haplotypes to present no variant. If both haplotypes have variants, we filter the allele;
- **Other**: alternative allele was excluded for other reasons;
- **High quality**: the final set of alternative alleles retained for further analysis.

Supplementary Table 1 shows the number of assembled alleles removed in each step.

#### Detecting split reads and hidden split reads supporting CNVs and expected support score

The Genome in a Bottle (GIAB) Consortium released a short-read sequencing dataset for HG002 using the Illumina HiSeq 2500 platform, and the New York Genome Center (NYGC) released a dataset for HG00512 sequenced with the Illumina NovaSeq 6000 platform. Using these datasets, we show that many CNVs in repetitive regions are not supported by split reads, but they are supported by hidden split reads.

For each CNV for which we obtained a high quality assembly, we identify the split reads and hidden split reads supporting it as follows. We align the alternative allele to the reference genome to calculate the precise coordinates of the breakpoints of the CNV, both on the alternative allele and on the reference genome. Then, we map the short reads on the alternative allele using BWA MEM [13], and we extract all reads overlapping a breakpoint. These reads should, in theory, support the existence of the CNV when mapped to the reference genome. Let us call $S$ the set of such reads.

We extract the reference sequence $R_{ref}$ between the start and the end of the CNV, plus 2000 bp of flanking sequences in each direction. Then, we create a second sequence $R_{cnv}$ by introducing the CNV in $R_{ref}$ (i.e., deleting the deleted part or inserting the inserted sequence). Finally, we map the reads in $S$ to both $R_{ref}$ and $R_{cnv}$: The reads that are clipped when aligned to $R_{ref}$ but not when aligned to $R_{cnv}$ are split reads supporting the CNV, and reads that are not clipped and have a better alignment score when aligned to $R_{cnv}$ than when aligned to $R_{ref}$ are hidden split reads supporting the CNV.

In order to calculate the expected score (ES) for a CNV, we fix a read length $l$, and sample from the alternative allele all of the possible reads of length $l$ containing the breakpoint(s). In other words, let $b_1$ and $b_2$ be the breakpoints of the CNV on the alternative allele $A$ and let $R = \{A[i..i+l] \mid i < b_1 < i+l \text{ or } i < b_2 < i+l\}$ be the set of all possible reads sequenced from the alternative allele that contain a breakpoint. Let $R'$ be the subset of reads in $R$ that aligns better to $R_{cnv}$ than to $R_{ref}$: then, $ES = \frac{|R'|}{|R|}$.

#### Realigning individual hidden split reads to detect potential CNVs

In order to show that hidden split reads must be used carefully, we randomly select 10,000 repetitive regions in hg38. For each tandem repeat region, we select all reads that do not align perfectly to the reference. For each read, we split it into two at every possible location, and locally align the two half reads independently to the repetitive region. We select the split that has the highest score (calculated as the alignment score of the left half of the read plus the alignment score of the right half of the read). Let $s_l, e_l$ be the start and end coordinates of the optimal alignment of the left half of the read, and $s_r, e_r$ be the start and end coordinates of the optimal alignment of the right half of the read. If $s_r > e_l$, we detect a deletion from $e_l$ to $s_r$; vice versa, if $s_r < e_l$, we detect a duplication from $s_r$ to $e_l$. Finally, we retain all CNVs $\geq$ 50bp.

#### Clustering split reads

We aim at clustering clipped reads and hidden split reads and, for each cluster of size at least three, generating a consensus sequence. Each consensus sequence represents a potential breakpoint of a CNV, and it is marked as right-clipped or left-clipped.

In the current algorithm, clipped reads and hidden split reads are not clustered together; in other words, a cluster cannot contain both clipped reads and hidden split reads. Left-clipped (resp. right-clipped) reads that are clipped at the same position (with a tolerance of 3 bp) are clustered together.

For hidden split reads, we heuristically determine whether a read is left or right-clipped as follows. We divide it into two halves, and count the number of differences with the reference (defined as number of mismatches plus the number of indels) for the left and the right half. If the left half has a higher number of differences than the right half, we mark the read as left-clipped; otherwise, we mark it as right-clipped. Left-clipped (resp. right-clipped) hidden split reads are clustered so that the overlap within any pair of reads within a cluster is at least half the read length.

The consensus generation proceeds identically whether the reads in the cluster are clipped or hidden split reads. For each cluster of size at least three, a consensus sequence is generated by casting a majority vote base by base [18]. Then, every read in the cluster is realigned against the consensus sequence: if the alignment contains any indel or the fraction of mismatches is greater than $\epsilon$ ($\epsilon$ is a user-defined value that represents the maximum expected fraction of sequencing errors in a read, 0.04 by default), the read is removed from the cluster. If less than three reads are

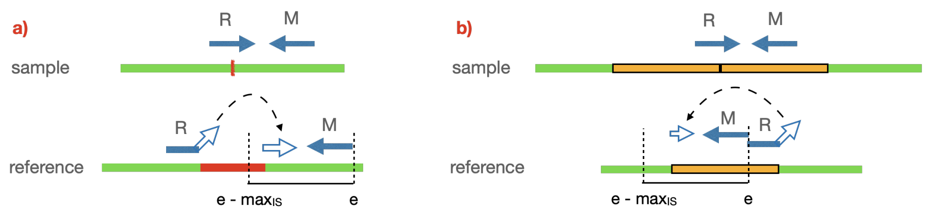


Figure S1: (a) An example of a right-clipped reads $R$ and its mate $M$. $R$ determines the left breakpoint of the deletion, and remapping the clipped sequence can determine the right breakpoint. By the definition of read pair, the right half of $R$ and $M$ must be within $max_{IS}$ bp of each other; therefore, we define as the acceptable range for the right breakpoint of the deletion to be $[e - max_{IS}..e]$, where $e$ is the endpoint of $M$. (b) For duplications the situation is similar, except that the $R$ determines the right breakpoint, and remapping the clipped sequence determines the left breakpoint. The acceptable range for the left breakpoint is therefore $[e - max_{IS}..e]$, where $e$ is the endpoint of $M$.

retained, the whole cluster is discarded. Otherwise, a new, final consensus sequence is generated from the retained reads. Left-clipped (right-clipped) reads will produce a left-clipped (right-clipped) consensus.

#### Finding a putative range for breakpoints

As mentioned in Section 2.2, the consensus sequence of a cluster of split reads represents a potential breakpoint of a CNV. Since a CNV has two breakpoints on the reference, we are interested in defining a range where to search for the second breakpoint. Fig. S1 shows how to calculate the range from a single right-clipped read. In order to calculate the range for a right-clipped cluster, we simply calculate the intersection of the ranges of the individual reads. The calculation of the range for left-clipped clusters is simply symmetrical.

#### Generating the junction sequence

Given a right-clipped consensus sequence $R[1..r]$ and a left-clipped consensus sequence $L[1..l]$, we generate a junction sequence as follows. Let $n$ be the largest integer such that the hamming distance between $R[r-n+1 : r]$ and $L[1 : n]$ is at most $n \cdot \epsilon$ (see Supp. Sec 4 for a definition of $\epsilon$). If $n \geq 15$, the junction sequence is formed by concatenating $R$ and $L[n + 1 : l]$; otherwise, no sequence is generated.

#### Extending a consensus sequence

Consensus sequences obtained by split and hidden split reads tend to be short, often not much longer than the length of a read. For this reason, when realigning them to repetitive regions we may obtain incorrect or ambiguous locations. By extending the consensus sequences using assembly we aim at improving the confidence of the realignment. This happens in two steps: first, a set of target reads are identified; second, the target reads are used to extend using an overlap-layout-consensus approach.

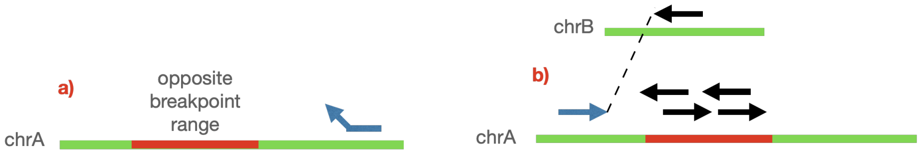


Figure S2: Detection of the target reads for the extension of a consensus sequence. (a) First, we identify the target region that contains the reads we will use to extend the consensus. In this case, since we are extending a left-clipped consensus to the left, the target region (in red) is the opposite breakpoint range of the consensus. (b) Next, we collect the reads within the target region (reads in black). Furthermore, an additional read is collected from an entirely different genomic location (described as chrB in the figure). This is because the location of its mate (in blue) suggests that the read might belong to the target region.

In order to detect the target reads, we first identify a target region where the desired reads are likely to be mapped. The target region is calculated as follows:

- If we are extending a right-clipped consensus to the left, the target sequence is the $maxIS$ bp region left-flanking the consensus;
- If we are extending a left-clipped consensus to the right, the target sequence is the $maxIS$ bp region right-flanking the consensus;
- If we are extending a right-clipped consensus to the right or a left-clipped consensus to the left, the target region is the opposite breakpoint range (see Section 2.2).

Then, all reads mapped within the target region are target reads. Furthermore, we search for pairs such that a read $R_1$ is aligned to the forward (resp. reverse) strand within $maxIS$ bp upstream (resp. downstream) of the target region and their mate $R_2$ is aligned to a different genomic location: $R_2$ is added to the target reads (Supplementary Fig. S2).

Next, we use the target reads to extend the consensus. A directed graph is built where each node is a read, and an edge connects two reads $R_1 \rightarrow R_2$ if a suffix of $R_1$ and a prefix of $R_2$ of at least read length/2 bp are identical. The consensus sequence is treated as a read. If the connected component containing the consensus sequence contains a cycle, we remove every edge that touches a node in the cycle. Then, we find the longest path either starting from or ending at the consensus sequence, depending on whether we are extending the consensus to the left or to the right.

#### Clustering discordant pairs

Given a set of discordant read pairs, we want to partition them into clusters so that all read pairs in a cluster support the same CNV. For this purpose, we use the algorithm used by SurVIndel [18]. Here we provide a brief, high-level description.

Each cluster is composed of two segments, a forward and a reverse segment. We define a merging operation between two segments as follows: given two segments $S_1 = (s_1, e_1)$ and $S_2 = (s_2, e_2)$, we create a new segment $S_m = (min(s_1, s_2), max(e_1, e_2))$. We then define a merging operation between two clusters by merging the two forward segments and the two reverse segments. We also define the distance between the two segments $S_1$ and $S_2$ is defined as $max(e_1, e_2) - min(s_1, s_2)$, and the distance between two clusters as the maximum distance between (a) the distance between the two forward segments and (b) the distance between the two reverse segments.

The algorithm proceeds as follows. Initially, each individual read pair forms a cluster, and its forward (resp. reverse) strand read is the forward (resp. reverse) segment of the cluster. Then, we keep merging the pair of clusters with the shortest distance, as long as the distance is less than $maxIS$. Intuitively, if a segment is longer than $maxIS$, it means that its cluster includes two reads that are more than $maxIS$ bp apart. Therefore, it is unlikely that such reads support the breakpoint of the same CNV.

#### Insertion size-based features

As mentioned in Section 2.2, CNVs will distort the insert size of read pairs. In particular, a read pair with a fragment size $L$ crossing a deletion of size $d$ will be mapped to the reference with an insert size of $L + d$; when $d$ is large, the read pair will be clearly discordant. However, when $d$ is small, determining that the pair supports the presence of a deletion is not as easy. Note that we define a read pair as discordant when the insert size is greater than $max_{IS} = \mu + 3\sigma$. Consider a read pair that contains a deletion, and assume its fragment size is $\mu$ bp. If the deletion is less than $3\sigma$ bp, the read pair will not be detected as discordant.

More precisely, define $min_{IS} = \mu - 3\sigma$ to be the minimum acceptable insert size for a read pair. Remember that $max_{IS}$ was defined similarly as the maximum acceptable insert size for a read pair. Then, when the deletion is of size greater than $max_{IS} - min_{IS} = 6\sigma$, we expect that all pairs containing the deletion can be identified as discordant, and we use a positive-to-negative ratio. Let $P$ and $N$ be the set of discordant and concordant (i.e., not discordant) pairs that contain the midpoint of the tested deletion. We compute the positive-to-negative ratio as $|P|/(|P| + |N|)$, and we reject the deletion if it is too low (lower than 0.25 by default).

However, this approach cannot be used for smaller deletions. This problem was tackled in SurVIndel [18] by analysing the distribution of all pairs that potentially contain the deletion. Although it may be impossible to determine whether a single pair supports the deletion or not, we expect the read pairs to have a larger insert size, on average, than a set of pairs that do not contain a deletion. Therefore, statistical tests are employed to determine whether a set of reads may contain a deletion.

Here, we calculate two features using two different tests. Both tests have two ingredients: a distribution $C$ of read pairs that do not contain a deletion, and a distribution $D$ of pairs that contain the candidate deletion that is being tested. $C$ is generated by choosing 1 million random locations in the genome and sampling the pairs that contain any of those locations. $D$ is the set of read pairs that contain the midpoint of the deletion. The first test determines a confidence interval for the size of the deletion, by computing a 99% confidence interval for the difference of means between $D$ and $C$, and tests whether the size of the predicted deletion falls within this range. The second tests employs a Kolmogorov-Smirnoff test to calculate a p-value estimating whether $C$ and $D$ are significantly different. The lower the p-value, the more likely $C$ and $D$ are different.

For the first test, the feature we obtain is 0 if the size of the deletion falls within the confidence interval, otherwise it is a positive number indicating how far outside the confidence interval the deletion falls. The second feature is the p-value of the KS-test.

The statistical tests are not currently applied to tandem duplications, because the distribution of reads pairs is far more complicated and challenging to use.

#### Comparing deletions and tandem duplications

When comparing two indels, we use a set of three parameters: the *max distance*, the *min overlap* and the *maximum length difference*. Two deletions or two tandem duplications are considered to match if:

1. The distance in bp between the two start coordinates and between the two end coordinates are less than or equal to *max distance*;
2. The fraction of the shortest indel that overlaps with the largest indel is at least *min overlap*;
3. The difference between the lengths of the two events is less than or equal to *maximum length difference*.

We use two sets of parameters, a *precise* (stricter) set and an *imprecise* (more permissive) set. When one or both the indels are marked as imprecise by the callers, we use the imprecise set, otherwise we use the precise set. For the precise set, we used (in order, max distance, min overlap, maximum length difference) 100 bp, 0.8, 100 bp. For the imprecise set, we used 500 bp, 0.5, 500 bp.

The same deletion or a tandem duplication in a tandem repetitive region may be represented using different coordinates. As an example, consider three genomic sequence A, B and C that appear as ABBBC in the reference and as ABBC in the sample (on B was deleted). Any of the copies of B in the reference may be marked as deleted and it would represent the same deletion. For this reason, we employ a repeat-aware comparison method. When two deletions or two tandem duplications fall within the same tandem repeat, we only check whether the lengths of the two events is less than or equal to maximum length difference.

As mentioned before, tandem duplications in the benchmark catalogues we used are represented as insertions, i.e., as an insertion site and an inserted sequence. In order to compare an insertion with a tandem duplication, we use a slightly modified method. The insertion site must be within *max distance* bp of either the start or the end of the duplication. Furthermore, we check whether the inserted sequence of the insertion and the duplicated sequence of the duplication match. Let $I[1..i]$ be the inserted sequence and $D[1..d]$ be the duplicated sequence in the reference. First, we find the number of times $D$ was duplicated as $n = \lceil \frac{d}{i} \rceil$, and create $D'$ by concatenating $D$ $n$ times. Then, we compute the local alignment between $I$ and $D'$ using a scoring scheme of +1, -4, -6, -1 (match, mismatch, gap opening, gap extension). If we alignment covers at least 80% of $I$, we accept that the insertion and the duplication match.

The comparisons performed for Fig. 7 were all performed using the imprecise parameters, to account for the possible shifting of the breakpoints due to clustering.

The code of the comparison algorithm is available at https://github.com/Mesh89/SurVClusterer.

#### Clustering deletions and tandem duplications

In order to cluster a set of indels, we use the algorithm described in [15]. First, the set of indels is represented as a compatibility graph, where every indel is represented as a vertex and the two indels are *compatible*, i.e., they potentially represent the same variant. We say that two indels are compatible if they match according to the comparison algorithm described in Supplementary Methods (we do not perform the repeat-aware comparison for efficiency reasons). After that, we heuristically compute the minimum clique cover (computing the optimal solution is not computationally feasible), and every clique is a cluster of indels. More details about the heuristic used can be found in [15]. The code of the clustering algorithm was published at https://github.com/Mesh89/SurVClusterer.

#### Other organisms genomic data

We downloaded Illumina paired-end and PacBio HiFi reads for seven Arabidopsis Thaliana: Alo-0 (ERR10084935 and SRR1946106), Cas-0 (ERR10084942 and SRR1946392), Cat-0 (ERR10084948 and SRR1946393), Cvi-0 (ERR10084078 and SRR1945758), Evs-0 (ERR10084952 and SRR1946405), Hom-4 (ERR10084958 and SRR1946144) and Hum-2 (ERR10084063 and SRR1946147). All of the samples were aligned to TAIR10, using BWA MEM for short reads and minimap2 for long reads.

For the Bos Taurus samples, we downloaded PacBio HiFi reads (ERR10378054 to ERR10378058) and two short read libraries, which we treated as independent (ERR10310239 and ERR10310240). We aligned the reads to the ARS-UCD1.3. For the Mus Musculus sample, we downloaded PacBio HiFi reads (SRR23686163) and short reads (SRR23690179), and aligned them to the GRCm39 reference genome. For Oryza sativa, we downloaded PacBio HiFi reads (SRR10238608 for MH63, SRR13280199 for ZS97) and short reads (SRR13124689 for MH63, SRR13124696 for ZS97), and aligned them to the IRGSP-1.0 reference genome.

Benchmark datasets for each sample were obtained by running Sniffles2 on the long read datasets.

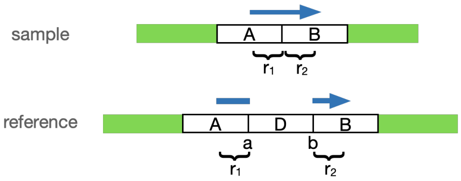


Figure S3: A region $D$, covering the reference from position $a$ to position $b$, is deleted. $A$ and $B$ are the left- and right-flanking regions of $D$, respectively. In the sample genome, $A$ and $B$ are adjacent. Therefore, a read $R$ of $r$-bp may be sequenced so that its $r_1$-bp long prefix is sequenced from $A$ and its $r_2$-bp long suffix is sequenced from $B$ (clearly, $r_1 + r_2 = r$). When $R$ is aligned to the reference, it will not completely align to any location. Rather, its first $r_1$-bp will align to the $r_1$-bp long suffix of $A$, and its last $r_2$-bp will align to the $r_2$-bp long prefix of $B$.

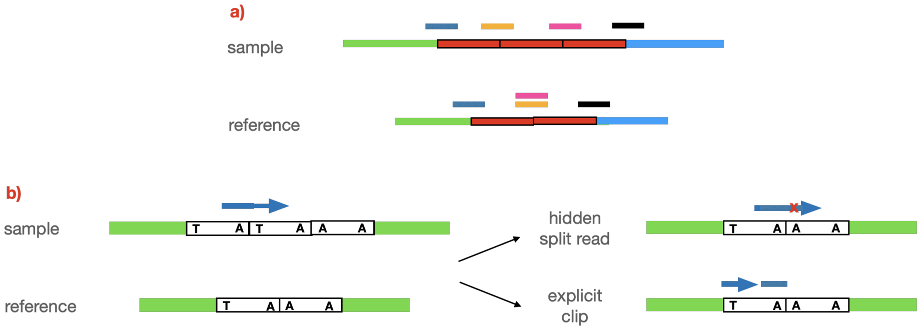


Figure S4: (a) Assume there is a tandem repeat with $n > 1$ repeat units in the reference, its repeat units are all identical, and its length is greater than the read length. If the same tandem repeat in the sample has $m > n$ repeat units, no split read will support the duplication. (b) Similarly to deletions, when copies of a tandem repeat are not identical, a duplication may generate hidden split reads.

### Supplementary Figures

### 4 Supplementary Data

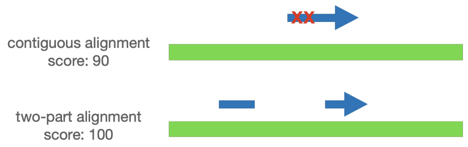


Figure S5: A read is aligned to the reference with two mismatches. Suppose we assign +1 to a match and -4 to a mismatch; its alignment score will be 90 (98 matches and 2 mismatches = 98 - 8 = 90). If we allow the read to split into two, and its two parts to align independently without penalty, we can align both parts without mismatches. Therefore, the score is 100 (100 matches). The HSR-score of this read is 100 - 90 = 10.

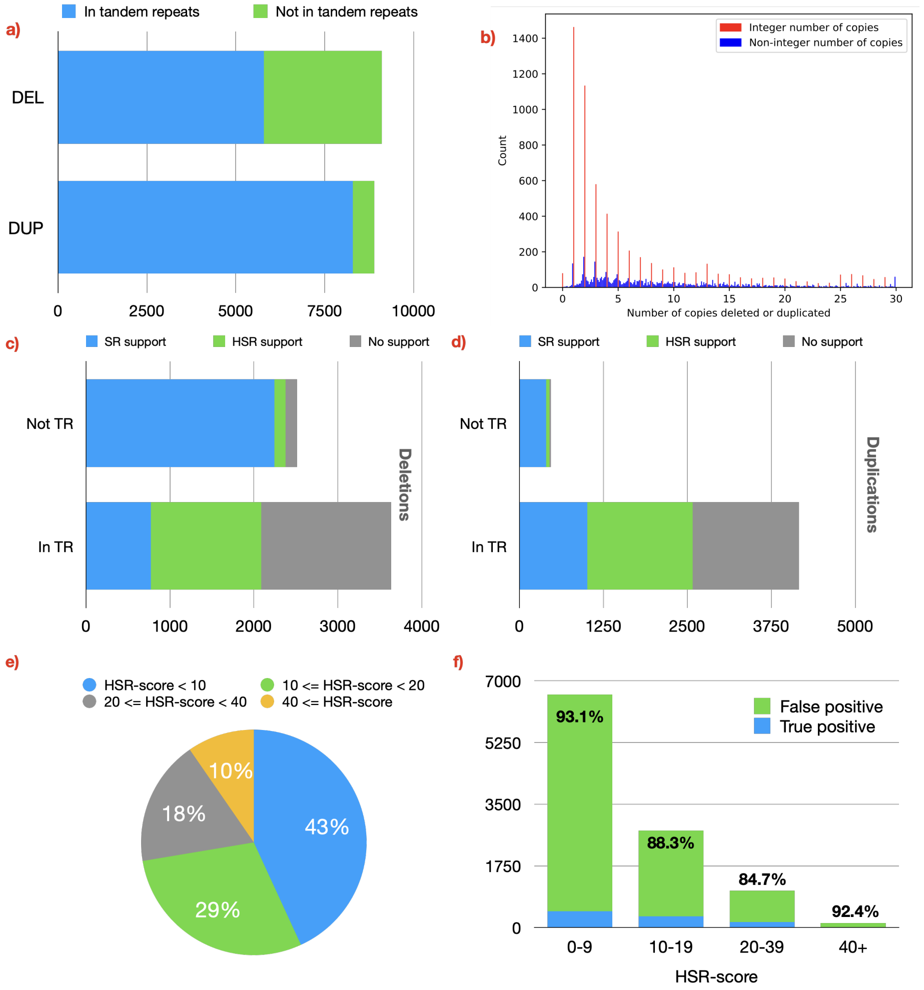


Figure S6: The impact of deletions deleting full repeat units and the importance of hidden split reads in detecting such deletions, studied on HG00512. (a) We consider a deletion to be contained in a tandem repeat if at least 90% of the deletion is covered by a repetitive region. Nearly two thirds of the deletions in this dataset are contained in a tandem repeat. (b) For every deletion that overlaps with a tandem repeat, we calculated the number of repeat units that it deletes by dividing the length of the deletion by the length of the repeat unit. Bars representing an integer number of copies (i.e., the length of the deletion is multiple of the length of the repeat unit) are coloured red. Most deletions delete an integer number of repeat units. (c,d) Percentages of deletions outside (c) and contained (d) in tandem repeats that are supported by split reads, by hidden split reads, and by neither. Hidden split reads have the potential to recover a significant number of deletions missed by regular split reads. (e) HSR-scores of the hidden split reads supporting the existing CNVs in HG00512. Most reads have low (<20) HSR-score. (f) We found all hidden split reads in 10,000 randomly sampled repetitive regions, and for each we tried to predict a CNV by finding the optimal split alignment. Most CNVs (>91%) detected this way were false positive. This shows that using hidden split reads to detect CNVs is challenging, and effective filters must be employed to distinguish real from false CNVs.

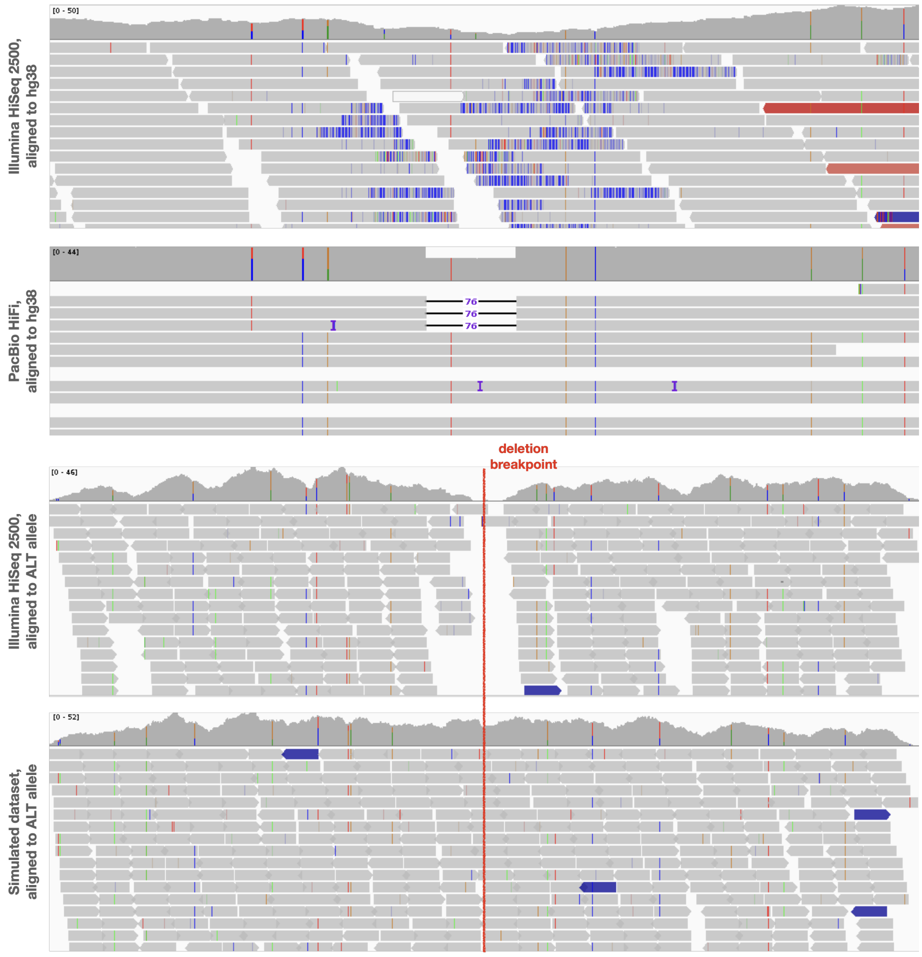


Figure S7: An example of unsupported deletion in the HGSVC2 catalogue for HG002, called chr10-729000-DEL-76. The deletion is clearly supported by the HiFi reads aligned to hg38, but the HiSeq 2500 reads in the region appear to be very noisy. When aligned to the alternative allele, we observe that only one good quality read (BWA-MEM AS$\geq$ 140, read length 150) contains the breakpoint. For this reason, despite having a high ES score (0.72), we cannot find reads supporting the deletion: for this reason, SurVIndel2 does not call it. However, in the simulated reads, plenty of correctly aligned reads contain the breakpoint. Unsurprisingly, SurVIndel2 can detect the deletion in the simulated dataset.

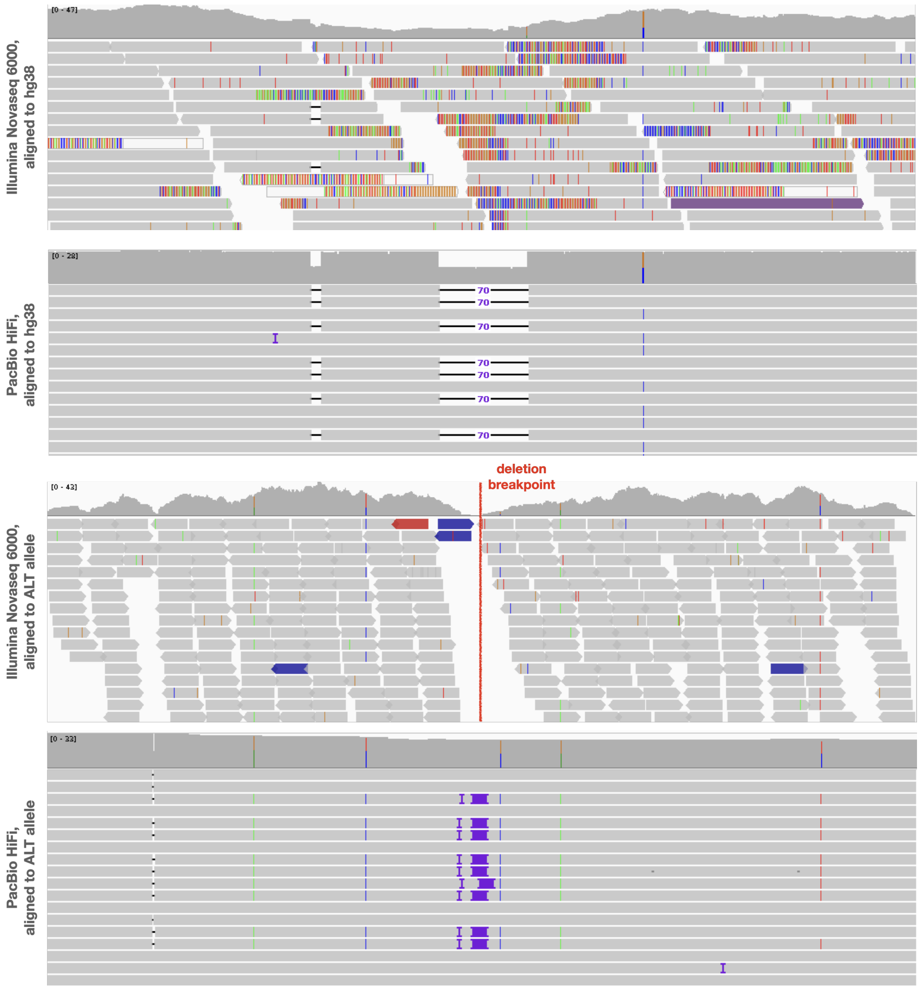


Figure S8: An example of unsupported deletion in the HGSVC2 catalogue for HG00512, called chr1-2262046-DEL-70. The deletion is clearly supported by the HiFi reads aligned to hg38, but the Illumina reads in the region appear to be very noisy. When aligned to the alternative allele, we observe that there are no good quality reads (BWA-MEM AS$\geq$ 140, read length 150) that contain the breakpoint. For this reason, despite having a high ES score (0.633), we cannot find any reads supporting the deletion. Many HiFi reads perfectly align the the alternative allele, which confirms that the alternative allele is correct.

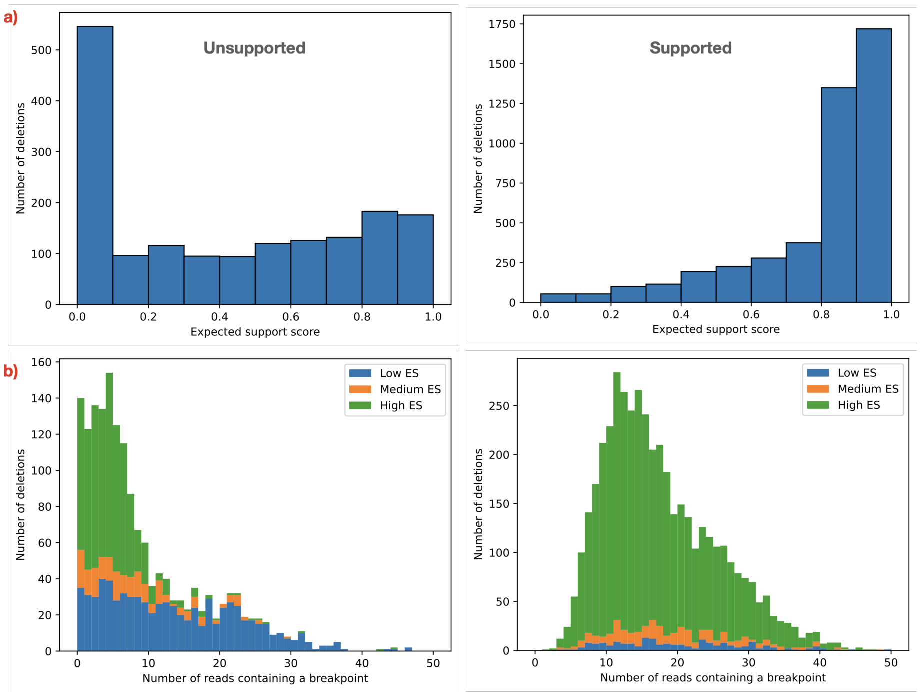


Figure S9: Factors that contribute to the absence of support for deletions in HG00512. The results confirm our findings on HG002. a) As for HG002, a large portion of unsupported deletions have low ES. However, 44% of the unsupported deletions have high EQ ($\geq 0.5$), which suggest that there is another reason the lack of support. b) Another factor is an absence of correctly sequenced reads containing the breakpoint of the deletions.

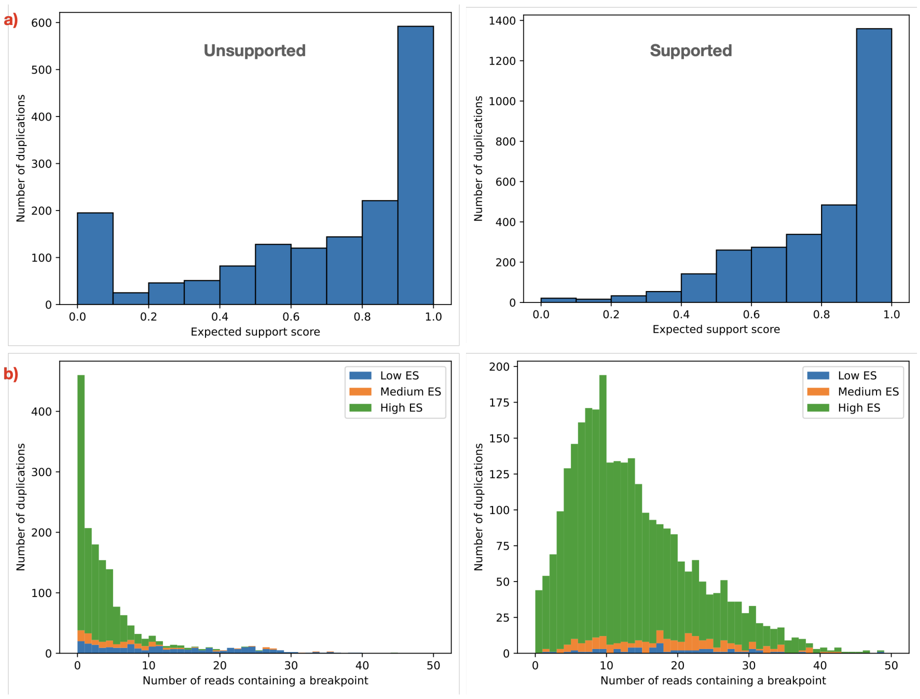


Figure S10: Factors that contribute to the absence of support for duplications in HG00512. The results confirm our findings on HG002. a) As for HG002, most unsupported deletions (75%) have high EQ ($\geq$ 0.5), which suggest that there is another reason the lack of support. b) The absence of correctly sequenced reads containing the breakpoints of the duplication is even more prevalent than in deletions.

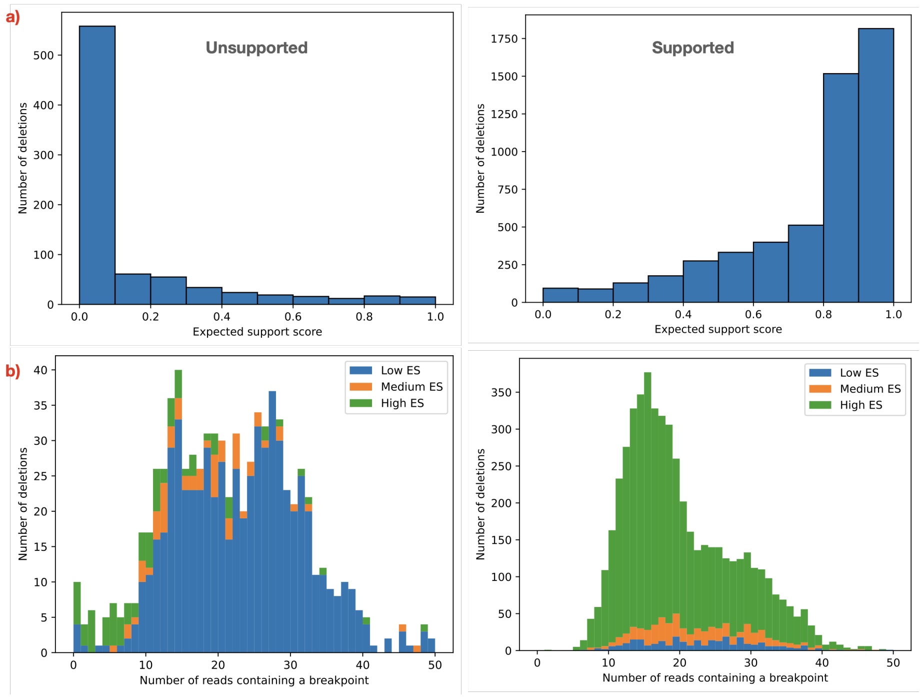


Figure S11: Factors that contribute to the absence of support for deletions, in a realistic synthetic dataset for HG002. a) As expected, the vast majority of unsupported deletions have low ES, while supported deletions have high ES. Unlike the real dataset (Fig. 9), nearly no unsupported deletions have high ES. b) The real dataset exhibited a large number of deletions that have high ES but no correctly sequenced reads containing their breakpoints. This phenomenon is absent from the synthetic dataset.

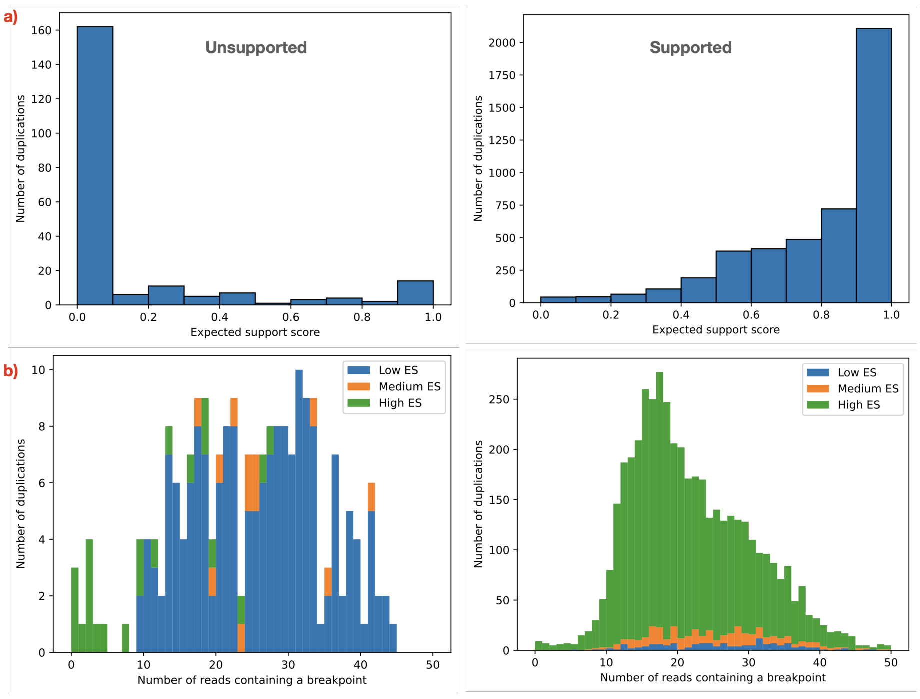


Figure S12: Factors that contribute to the absence of support for duplications, in a realistic synthetic dataset for HG002. a) As expected, the vast majority of unsupported duplications have low ES, while supported duplications have high ES. This is in stark contrast to the real dataset, where the majority of unsupported duplications had high ES (Fig. 10). b) The real dataset exhibited a large number of duplications that have high ES but no correctly sequenced reads containing their breakpoints. This phenomenon is absent from the synthetic dataset.

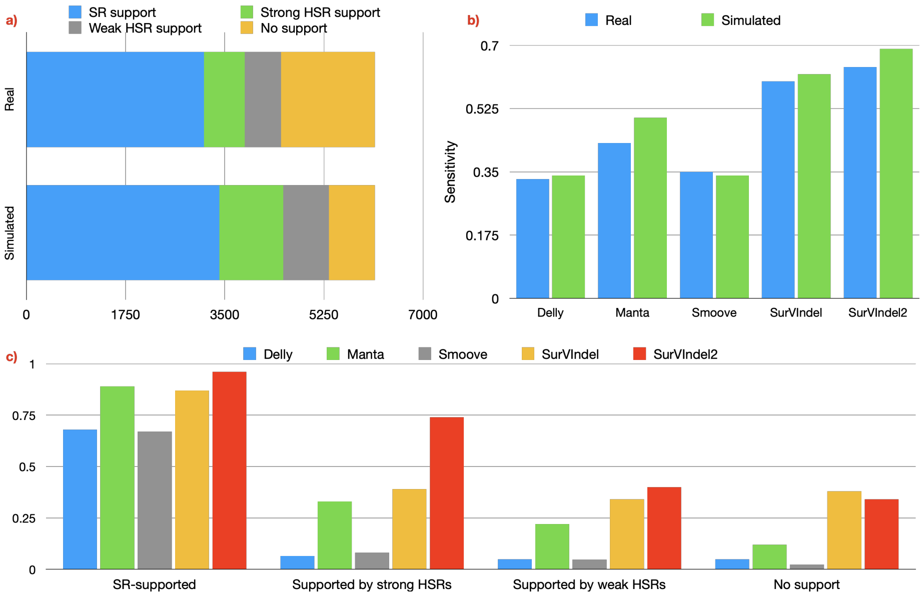


Figure S13: Support for deletions in the synthetic versus the real dataset. a) The simulated dataset has less unsupported deletions. b) The sensitivity of some methods is increased. This is particularly true for Manta (increase from 0.43 to 0.5) and SurVIndel2 (increase from 0.64 to 0.69). c) However, when stratified by evidence available, the sensitivity of the methods for each category are very similar between the real (Fig. 4c) and the synthetic datasets. This, combined with Supplementary Fig. S11 shows that the increase in sensitivity is due to less unsupported deletions.

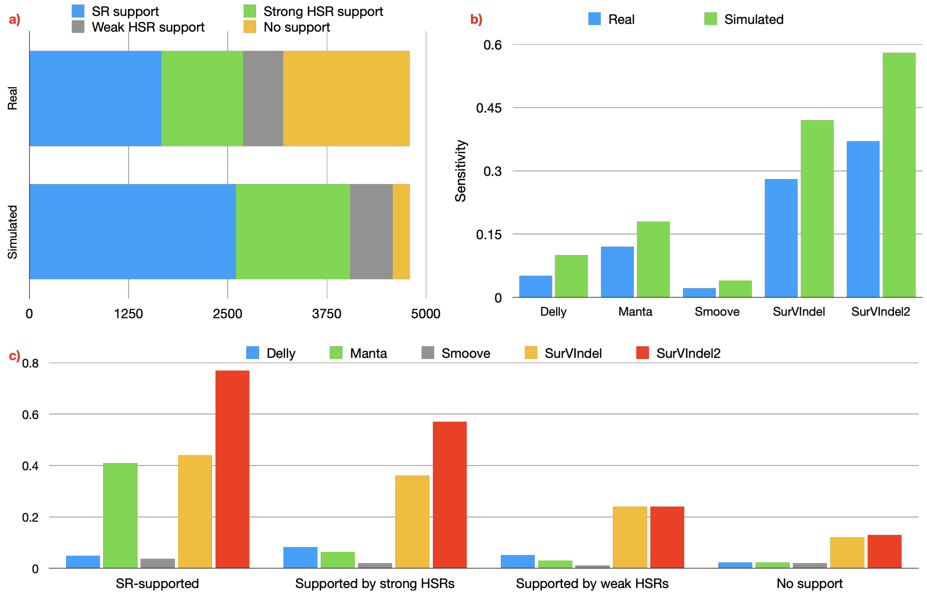


Figure S14: Support for tandem duplications in the synthetic versus the real dataset. a) The simulated dataset has far less unsupported duplications. b) The sensitivity of all the methods is considerably increased. Particularly notable is the performance of SurVIndel2 improves from 0.37 to 0.58. c) Once again, when stratified by evidence available, the sensitivity of the methods for each category are very similar between the real (Fig. 4d) and the synthetic datasets. This, combined with Supplementary Fig. S12 shows that the methods are severely limited because the regions containing the tandem duplications are not sequenced correctly.

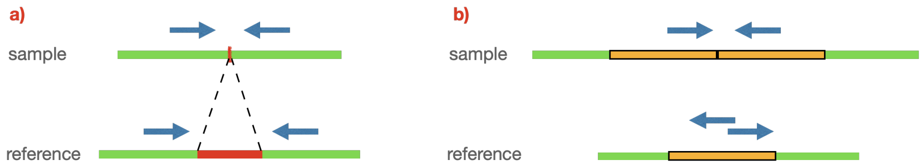


Figure S15: (a) Consider a read pair s.t. one read was sequenced upstream and one downstream of the deletion breakpoint, sequenced from a fragment of size $L$. After mapping the pair to the reference, its insert size will be $L + d$, where $d$ is the size of the deletion. If the deletion is large enough, the insert size will also be abnormally large, when compared to the insert size distribution of the library. (b) When a duplication is large, reads sequenced from different copies of the duplicated sequence may show the wrong relative orientation when aligned to the reference.

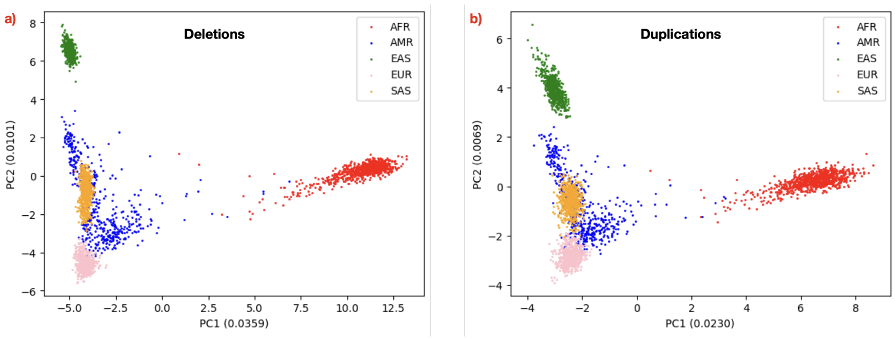


Figure S16: PCA based on deletions (a) and tandem duplications (b). Both event types are able to clearly separate the superpopulations.

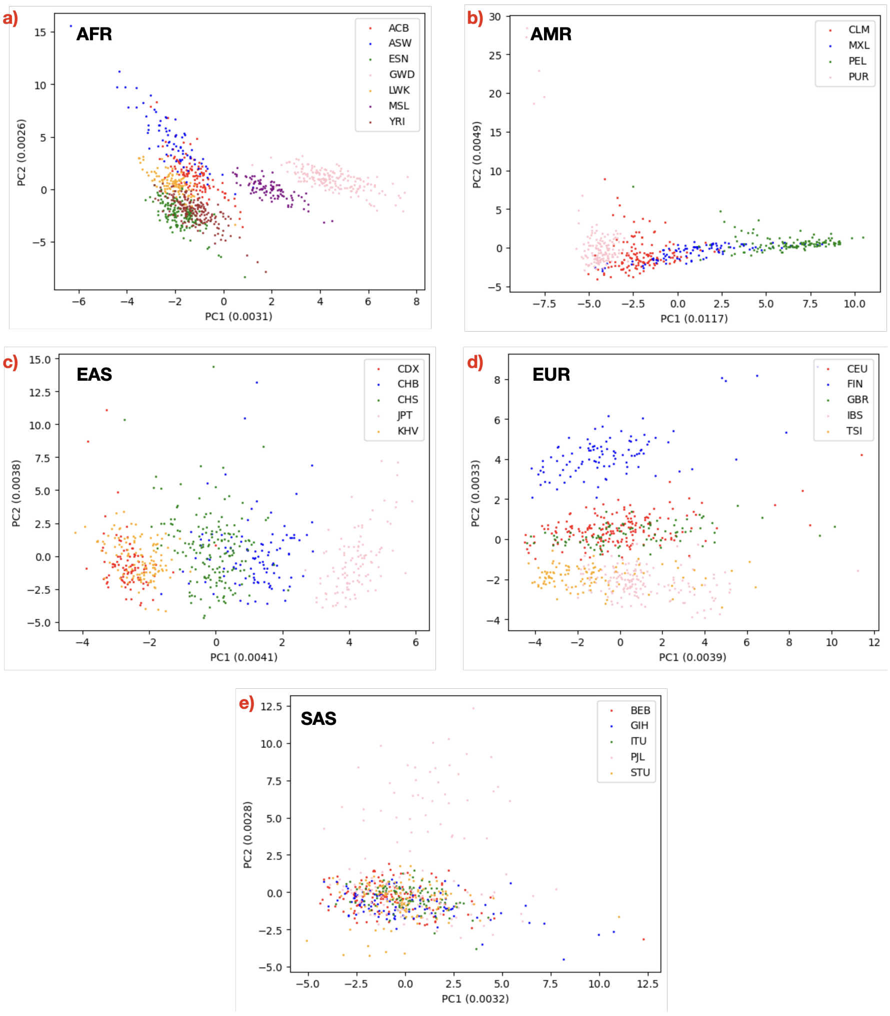


Figure S17: PCA based on both deletions and duplications was able to segregate each superpopulation into subpopulations. (a) For Africans (AFR), two subpopulations are clearly separated from the others: Mende in Sierra Leone (MSL) and Gambian in Western Division (GWD). Americans (AMR) subpopulations are segregated on PC1, while PC2 is dominated by heterogeneity among Puerto Ricans (PUR). (c) East Asians (EAS) form three major clusters: (1) Vietnamese and Dai Chinese, (2) Han Chinese from Beijing with Southern Han Chinese and (3) Japanese. Some degree of separation between Han Chinese from Beijing and Southern Han Chinese is also observed. Similarly (d), Europeans (EUR) are also clustered into three clusters: (1) Finnish, (2) British and Utah residents with Northern and Western European ancestry and (3) Italians from Tuscany and Spanish. (e) South Asians (SAS) structure is much less obvious than other superpopulations. Furthermore, there appears to be high heterogeneity among Punjabi from Lahore (PJL). These results agree what we previously observed from insertions [17]

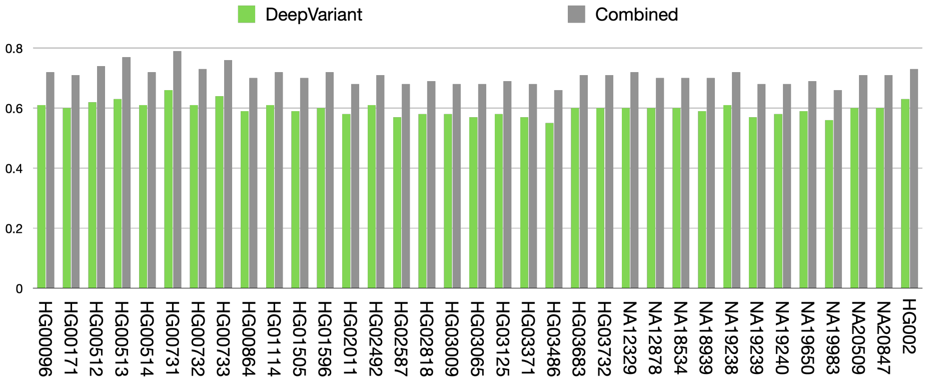


Figure S18: Sensitivity of DeepVariant and DeepVariant plus SurVIndel2 on the 35 HGSVC2 samples, for deletions between 30 and 50 bp.

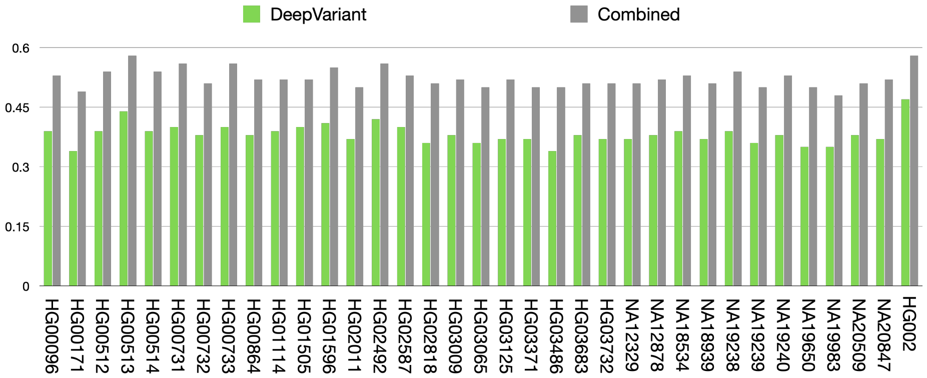


Figure S19: Sensitivity of DeepVariant and DeepVariant plus SurVIndel2 on the 35 HGSVC2 samples, for insertions between 30 and 50 bp.
